# Genome-scale modeling specifies the metabolic capabilities of *Rhizophagus irregularis*

**DOI:** 10.1101/2021.10.07.463607

**Authors:** Philipp Wendering, Zoran Nikoloski

**Affiliations:** Bioinformatics, Institute of Biochemistry and Biology, University of Potsdam, 14476 Potsdam, Germany; Systems Biology and Mathematical Modeling, Max Planck Institute of Molecular Plant Physiology, 14476 Potsdam, Germany

## Abstract

*Rhizophagus irregularis* is one of the most extensively studied arbuscular mycorrhizal fungi (AMF) that forms symbioses with and improves the performance of many crops. Lack of transformation protocol for *R. irregularis* renders it challenging to investigate molecular mechanisms that shape the physiology and interactions of this AMF with plants. Here we used all published genomics, transcriptomics, and metabolomics resources to gain insights in the metabolic functionalities of *R. irregularis* by reconstructing its high-quality genome-scale metabolic network that considers enzyme constraints. Extensive validation tests with the enzyme-constrained metabolic model demonstrated that it can be used to: (1) accurately predict increased growth of *R. irregularis* on myristate with minimal medium; (2) integrate enzyme abundances and carbon source concentrations that yield growth predictions with high and significant Spearman correlation (*ρ_S_* = 0.74) to measured hyphal dry weight; and (3) simulated growth rate increases with tighter association of this AMF with the host plant across three fungal structures. Based on the validated model and system-level analyses that integrate data from transcriptomics studies, we predicted that differences in flux distributions between intraradical mycelium and arbuscles are linked to changes in amino acid and cofactor biosynthesis. Therefore, our results demonstrated that the enzyme-constrained metabolic model can be employed to pinpoint mechanisms driving developmental and physiological responses of *R. irregularis* to different environmental cues. In conclusion, this model can serve as a template for other AMF and paves the way to identify metabolic engineering strategies to modulate fungal metabolic traits that directly affect plant performance.

**Importance:** Mounting evidence points at the benefits of the symbiotic interactions between the arbuscular mycorrhiza fungus *Rhizophagus irregularis* and crops; yet, the molecular mechanisms underlying the physiological responses of this fungus to different host plants and environments remain largely unknown. We present a manually curated, enzyme-constrained genome-scale metabolic model of *R. irregularis* that can accurately predict experimentally observed phenotypes. We show that this high-quality model provides an entry point into better understanding the metabolic and physiological responses of this fungus to changing environments due to the availability of different nutrients. The model can be used to design metabolic engineering strategies to tailor *R. irregularis* metabolism towards improving the performance of host plants.

## Introduction

More than two thirds of all land plants are involved in symbiotic relationships with arbuscular mycorrhizal fungi (AMF) (1). AMF are members of a monophyletic group within the early diverging fungi. Arbuscular mycorrhizal symbiosis is established by fungal hyphae entering cortical root cells of the host plant to form subcellular structures, termed arbuscles (ARB), where nutrients are exchanged between the symbiotic partners (2, 3). *Rhizophagus irregularis* (previously wrongly ascribed to *Glomus intraradices* (4)) is one of the most extensively studied AMF, shown to form symbioses with a variety of agriculturally relevant plants. Soil inoculation with *R. irregularis* leads to improved overall plant growth (5–8), fruit quality (9, 10), and yield (11–14). Further, *R. irregularis* confers robustness against multiple abiotic stress conditions (15–22). These qualities thus make it a valuable contributor to plant fitness, which is widely exploited for plant cultivation.

Spores of *R. irregularis* grow into a network of coenocytic hyphae, which can be separated into three major structures: the extraradical mycelium (ERM), the intraradical mycelium (IRM), and ARB (2). The ERM is comprised of hyphae located in soil, whereas hyphae of the two apoplastic structures, IRM and ARB, grow between or penetrate cortical root cells. *R. irregularis* mainly provides inorganic phosphate (Pi) and nitrogen (N) to the host plant as its extensive hyphae network bridges the nutrient depletion zone surrounding the roots (23–27); in return, it receives carbohydrates and lipids from the host plant (28–35). Pi is one of the key nutrients that limits plant growth, and under Pi-limiting conditions, most plants rely on additional Pi supplied by a fungal symbiotic partner (3). To this end, the external hyphae of the fungus either take up Pi directly from the soil or obtain it from hydrolysis of complex organic phosphates, such as phytate (36). According to the current evidence in *R. irregularis*, assimilated Pi is polymerized into polyphosphate (PolyP), which is translocated through the ERM towards IRM (27). Finally, Pi is released from arbuscles into the periarbuscular space. Several Pi transporters have been identified in *R. irregularis* which could be involved in Pi translocation from fungus to plant (26, 37, 38).

Moreover, N is another key nutrient for plant growth, comprising up to 5% of their dry weight. However, the availability of N sources to the plant is restricted due to the limited range of roots and its inhomogeneous distribution in soil. Hence, many plants depend on interactions with microbes which can provide additional nitrogen assimilated from the surrounding soil (39). *R. irregularis* takes up N in the form of ammonia (NH_4_^+^) and nitrate (NO_3_^-^) as well as amino acids and small peptides via designated transporters. Three NH_4_^+^ transporters, GintAMT1-3, and a NO_3_^-^ transporter, GiNT, have been identified in *R. irregularis* (40–43). Intracellular NH_4_^+^ is then used to synthesize L-arginine from L-glutamate (25, 43). Arginine is assumed to be the major transport form of nitrogen from the ERM to IRM, where it is catabolized to NH_4_^+^ and excreted into the periarbuscular space (3, 43).

The fungus, in turn, is dependent on carbohydrates and lipids obtained from the plant host. Multiple sugar transporters have been found, which are likely involved in hexose transfer from the host plant to *R. irregularis* (31, 44). However, the sugars obtained from the plant are not sufficient for the fungus to complete its life cycle (i.e. formation of fertile spores). *R. irregularis* cannot synthesize fatty acids with chain length greater than eight due to the absence of the fatty acid synthase (FASI), and thus depends on fatty acids provided by the host plant (32, 33, 35, 45, 46). Most likely, lipid is transported as 2-monopalmitin; however, it has also been shown that *R. irregularis* is able to grow on myristate (47). These findings have been exploited to develop an axenic culture medium on which the obligate biotroph can grow up to the production of fertile spores (48).

The availability of an assembled genome for *R. irregularis* (49–52) largely facilitated the characterization of transporters and its lipid metabolism (45, 53), allowing us to draw conclusions about the metabolic capabilities of the obligate biotrophic fungus. Multiple studies performed gene expression profiling under various conditions, facilitating a deeper understanding of the *R. irregularis* metabolism and arbuscular mycorrhiza (5, 54–56). An annotated genome of an organism is also the basis for the generation of genome-scale metabolic models (GEMs) that offer the means to *in silico* probe the functional capabilities and physiological responses of the organism (57). GEMs have already been developed to analyze the interaction of a N-fixing bacterium *Sinorhizobium meliloti* and its host plant *Medicago truncatula* (58, 59). As a result, important features of the N exchange and co-dependent growth were revealed, leading to a better understanding of this symbiotic relationship. Such analyses for *R. irregularis* cannot be performed due to the lack of a high-quality GEM for this organism.

Availability of a GEM for *R. irregularis* can be particularly useful to dissect mechanisms underlying arbuscular mycorrhiza and to predict fungal nutrient conversions and exchange, directly affecting growth of the host plant. Here, we present a compartmentalized enzyme-constrained GEM for *R. irregularis*, termed iRi1574, which allows the integration and prediction of transcript and protein abundances for different growth scenarios. We then used the enzyme-constrained GEM of *R. irregularis* to predict protein abundances across four carbohydrate sources and three feeding concentrations; we also examined the predictions of growth and pathways that affect this complex phenotype based on experimental measurements of hyphal dry weight and protein content from Hildebrandt et al. (60). We show that the enzyme-constrained iRi1574 model results in predictions that correlate well with experimentally measured dry weight (as well as calculated growth rates) and allows us to probe the flux redistributions across three fungal structures using re-analysed published gene expression data (5). Thus, we lay the foundation for further in-depth analysis of *R. irregularis* metabolism, hypothesis testing regarding mechanism essential for arbuscular mycorrhiza, and metabolic engineering of this fungus to improve the effect on agriculturally relevant plant traits.

## Results and Discussion

### Reconstruction of the genome-scale metabolic model of *R. irregularis*

Our first contribution is the generation of a GEM for *R. irregularis* encompassing all enzymatic functions annotated for this agronomically relevant fungus. The metabolic model can be used in combination with computational approaches from the constraint-based modelling framework to predict variety of metabolic phenotypes, including growth, in different scenarios (61, 62). The genome of *R. irregularis* (49, 51) was used as a starting point for the generation of the GEM using the KBase fungal reconstruction pipeline (63). The resulting draft model was first translated to a common namespace, based on augmenting a database of biochemical reactions, ModelSEED (34), since there were reaction and metabolite identifiers from published fungal models without cross references. We then added 198 transport reactions from the *Saccharomyces cerevisiae* iMM904 GEM (64) to improve the network connectivity (Suppl. Tab 8). We further expanded the list of reactions based on literature evidence for *R. irregularis* (Suppl. Tab 1). After these steps, the model was manually curated to ensure mass- and charge balancing. Finally, stoichiometrically balanced cycles were removed from the model to avoid simulations in which growth without available carbon source is possible (Supp. Note 1).

The manually curated GEM of *R. irregularis*, named iRi1574, consists of 1286 metabolites and 1574 reactions in eight sub-cellular compartments, i.e. the cytosol, mitochondrion, peroxisome, the Golgi apparatus, Endoplasmic Reticulum, nucleus, vacuole, and an extracellular compartment. In total, 687 enzyme-coding genes are associated with 1054 (67%) reactions via gene-protein-reaction (GPR) rules (Fig. 1A). Further, we cross-referenced both metabolites and reactions to commonly used biochemical databases to increase the comparability to other GEMs and to facilitate its future usage. A published cost-efficient medium, that is used in dual-compartment culture systems and includes: glycine, myo-Inositol, pyridoxine hydrochloride, thiamine hydrochloride, nicotinic acid, and essential minerals, is the default medium for simulations (65). The dependence of the growth of *R. irregularis* on lipid transferred from the host (most likely 16:0 β-monoacylglycerol (32, 33)) was modelled by adding an exchange reaction for palmitate, which is added to the default medium.

**Figure 1.**
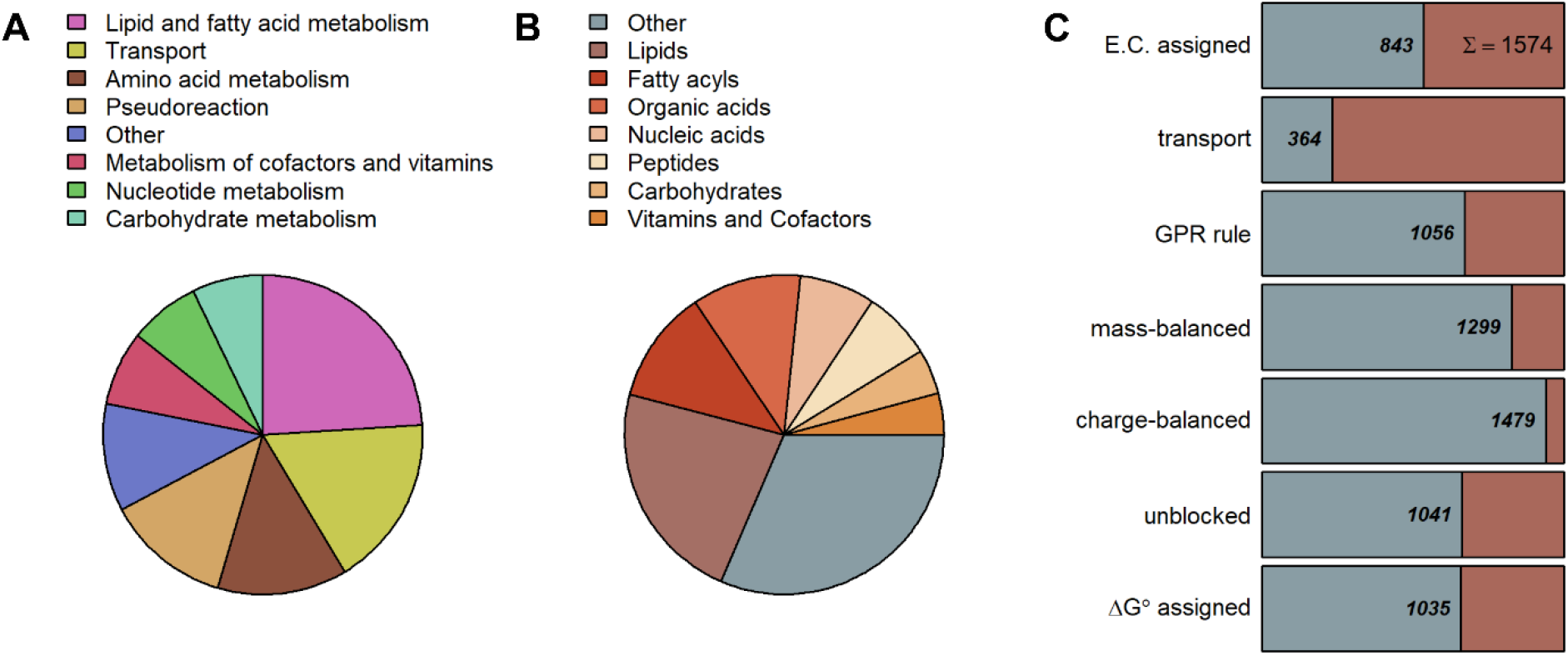
Properties of the *R. irregularis* genome-scale metabolic model iRi1574. (**A**) The iRi1574 includes 13 metabolic subsystems, primarily defined by KEGG pathways with manual refinement. The pie chart illustrates the percentage of reactions participating in these metabolic subsystems. (**B**) metabolite classification using KEGG BRITE with manual refinement with help of the ChEBI ontology. (**C**) binary classification of reactions based on eight criteria, including: assignment of Enzyme Classification (E.C.) number, involvement in transport, association to genes via GPR rules, mass- and charge-balancing, available value for standard Gibbs free energy and catalytic constants of associated proteins, and ability to support steady-state flux.

Altogether, the iRi1574 model includes 13 metabolic subsystems (Fig. 1B). In total, 24% and 13% of reactions take part in lipid and amino acid metabolism subsystems, respectively, which dominate the reconstruction (Fig. 1B). To model the lipid metabolism of *R. irregularis*, we relied on the gene annotations supported by literature (45, 66). Moreover, to incorporate experimentally measured lipid abundances (45, 67) into the biomass reaction, we used the SLIMEr method (68), whereby specific lipid species are split into their fatty acyl chains and backbone which are then combined using respective pseudoreactions. Hence, the number of lipid-related reactions and pseudoreactions is high compared to that of the remaining 11 metabolic subsystems. Based on literature evidence, we further added reactions that allow the production of ethylen (69), short-chain lipochitooligosaccharides (LCO) (70, 71) and vacuolar polyphosphate (72, 73). The respective end-products of these reactions are exported via sink reactions. Moreover, we added extracellular sink reactions for organic phosphate and ammonia as these molecules are known to be transported from the fungus to the host plant.

As only a small proportion of metabolites is annotated by the KEGG BRITE hierarchy (74), we used the ChEBI metabolite ontology (75) to structurally classify the considered metabolites. Due to the large number of reactions from lipid metabolism included in iRi1574, the proportions of lipids and fatty acyls are high (34%), followed by peptides/amino acids, organic-, and nucleic acids (Fig. 1C). The class of ‘Other’ metabolites is dominated by carbonyl compounds, heterocyclic compounds and phospho sugars. Quality assessment tests with the iRi1574 model were performed employing the MEMOTE test suite (76), yielding an overall score of 72% (Suppl. Note 1, Suppl. File 2).

### Comparison of iRi1574 to other fungal models

As *R. irregularis* is phylogenetically distant from other fungi for which GEMs have been published, we next asked whether the phylogenetic relationship among these fungi is represented in the enzyme sets included in the respective GEMs. To this end, we assigned pathway information to the reaction in nine fungal models according to the classification contained in the YeastGEM v8.3.5 (77) model (Suppl. Tab 2). To determine the overall similarity between two fungal models, we determined the overlap in E.C. numbers per subsystem by using the Jaccard Index (JI). We observed that, in comparison to the nine compared fungal models, the iRi1574 showed the lowest JI, i.e. lowest overlap of E.C. numbers, for fatty acid metabolism (including synthesis and elongation), thiamine metabolism, glycerolopid, and nicotinate and nicotinamide metabolism (Figure S1). In contrast, the largest overlap was found for the pentose phosphate pathway, one carbon pool by folate, pantothenate and CoA biosynthesis as well as amino sugar and nucleotide sugar metabolism, to name a few (Figure S1). Further, we identified that are some fungal GEMs showing differences in comparison to iRi1574 with respect to particular metabolic subsystems; for instance, the model of *N. crassa* showed particular differences in the tricarboxylic acid (TCA) cycle and pyruvate metabolism, the model of *A. terreus* displays particular differences in purine metabolism, steroid biosynthesis, sphingolipid, and pyrimidine metabolism, and lipid metabolism, while the model of *P. chrysogenum* differed in sphingolipid metabolism and fatty acid elongation (Figure S1).

The previous comparison between the fungal models was conducted only with respect to overlap of E.C. numbers present in particular metabolic subsystems, and does not point at differences in the activity of these pathways and their contribution to the physiology of the modelled fungi. To address this issue, we employed Flux Balance Analysis (FBA)(78, 79), that facilitates simulation of growth at steady state in each of the fungal models by optimizing of the flux, *ν_bio_*, through a biomass reaction that integrates the biomass precursors. This results in a linear optimization problem that imposes metabolic steady state and physiologically relevant bounds on reaction fluxes, i.e.

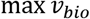

s.t.

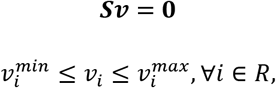

with ***S*** representing the stoichiometric matrix, including the molarity with which each substrate and product enter a reaction of the metabolic model, ***ν*** stands for the flux distribution, and *R* denoting the set of reactions in the model. Since it is well-known that there are, often, multiple steady-state flux distributions, ***ν***, that achieve the same growth (80), to characterize the activity of a metabolic subsystem, we next determined the minimum and maximum values that the sum of fluxes of the reactions participating in a given subsystem attain at optimal growth (see Methods). Similarly, we determined the sums of fluxes from parsimonious FBA (pFBA) for each of the subsystems (see Suppl. Methods).

Following this analysis, we observed that the ranges between the maximal and minimal sums of fluxes are largely overlapping and are of similar widths across most of the compared models (Figure S2). Interestingly, the model for *P. Chrysogenum*, iAL1006, and the iRi1574 model showed narrower ranges compared to the remaining models, except for fatty acid metabolism. Moreover, we observed that the maximum sum of fluxes is similar across all fungal models (coefficient of variation, 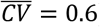), while minimal sums and sums from pFBA fluxes showed larger differences (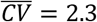 and 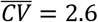). This suggests that these pathways are of differential importance for the models, since the minimal sum of fluxes provides an indication of how much flux must at least pass through these reactions to guarantee optimal growth. In conclusion, we find differences in both E.C. number overlap as well as in the pathway activities between the iRi1574 and other fungal models, indicating that iRi1574 is both structurally and functionally distinct from other fungal GEMs.

### iRi1574 can predict phenotypes of *R. irregularis* in line with experimental observations

We employed the assembled GEM to predict physiological traits for which there exists sufficient evidence and, thus, can be used to validate the performance of the model. A first question is how many of the reactions in the assembled model can carry flux. For these simulations, the M-medium(65, 81) was used, which was enriched with palmitate, D-glucose, and D-fructose, assumed to be supplied by the plant (Table S11). Using this default medium, 658 (42%) reactions were blocked (i.e. could not carry flux in any steady state supported by the model) of which 105 are transport reactions for extracellular metabolites. This is in line with the percentage of blocked reactions in the fungal models used in the comparison above (from 11.9% in iJL1454 to 49.9% in iRL766).

An important characteristic of the symbiotic relationships formed by *R. irregularis* is its dependence on association with the plant host to ultimately form fertile spores (46). According to recent findings, lipids are supplied by the plant symbiont, as *R. irregularis* does not possess the required enzyme set for *de novo* synthesis of long-chain fatty acids from hexoses (35, 45). More specifically, 2-monopalmitate was proposed as a likely candidate for the lipid exchange from plant to fungus (32, 33). Concordantly, axenic growth of this fungus is only possible when fatty acids are supplied in the medium (47, 82). Hence, the default medium used in the study includes palmitate as a lipid source. Indeed, simulations in which palmitate influx is blocked lead to no growth with or without consideration of other carbon sources.

It has been shown that *R. irregularis* is able to utilize additional carbon sources (30, 47, 83). The ability of the model to reproduce these finding was assessed by growth simulations on single carbon sources in the default medium, while restricting the uptake of palmitate to a minimal value that still guarantees optimal growth (8.46 *mmol gDW*^−1^*h*^−1^). As a result, we simulated growth on 11 carbon sources by using FBA (see above), resulting in different growth rates (Figure S3). Here, we observed the highest growth rates for trehalose, followed by D-glucose, D-fructose, melibiose, and raffinose. The observed high growth rate with trehalose as a carbon source is not surprising, given that it directly enters the biomass reaction. The equal growth rates obtained upon adding D-glucose, D-fructose, raffinose, and melibiose indicated that D-glucose, D-fructose, as well as D-galactose as a breakdown product from raffinose can be used with equal efficiency. The efficiency of the remaining carbon sources differed due to the differences in their breakdown pathway and additional modifications (e.g. phosphorylation, reduction).

Moreover, it has been reported that the addition of myristate to the medium leads to enhanced growth of *R. irregularis* (47). We found that optimum growth is, as expected, associated with a fixed value of palmitate influx of 8.46 *mmol gDW*^−1^*h*^−1^. Further, myristate is not utilized if additional carbon sources are unlimitedly available in the medium, which is in contrast to the experimental findings of Sugiura and co-workers, who found that the addition of myristate lead to an increment in growth irrespective an additional carbon source (47). Therefore, we asked if the reduced growth, due to the suboptimal scenario of fixing the palmitate influx to 10% of the minimum at optimal growth, can be complemented by adding myristate. Indeed, the model predicted that growth increased by 1.5% in comparison to the suboptimal scenario. When additional carbon sources (i.e. D-glucose, D-fructose, glycine, and myo-inositol) are allowed, with uptake rates restricted to their minimal fluxes at optimal growth, this increase in growth amounts to 9.7% (see below for the predictions from the enzyme-constrained model).

Another important transport process described for this symbiosis is the transport of Pi from fungus to the host plant (38). We found that the reconstructed model predicts export of Pi at optimal growth (Suppl. Tab 3 for FVA), in line with evidence (38). These results corroborate the quality of the functionally relevant predictions based on the developed iRi1574 model.

### Protein usage with different carbon sources

Enzyme-constrained GEMs have been developed for *S. cerevisiae* and *E. coli* (68, 84, 85), demonstrating improved prediction of metabolic phenotypes in contrast to the classical FBA-based models. In enzyme-constrained GEMs, the fluxes of reactions are bounded by the catalytic efficiency (*k_cat_* parameters) and the abundance of the respective enzyme(s) (86); these models also include constraints on the total enzyme content, borrowing from the initial idea proposed in FBA with molecular crowding (87, 88). An enzyme-constrained GEM can be used to predict not only growth, but also distribution of the total enzyme content across the different reactions and pathways. To generate an enzyme-constrained GEM for *R. irregularis*, we made use of 1214 *k_cat_* parameters, of which 430 were measured for fungi, that covered 57.4% of reactions included in the model (with all irreversible reactions, see Methods). We then employed an extension of MOMENT (84), a constraint-based approach that facilitates the integration and prediction of protein abundance by considering data on the *k_cat_* values. In addition to a molecular crowding constraint (Eq. 5, Methods) (84, 87, 88) similar to GECKO (68), we introduced a constraint to model enzyme promiscuity (Eq. 4), resulting in the extended method we refer to as eMOMENT. Missing turnover numbers were accounted for by assigning the median of the assigned *k_cat_* values.

Here, we first revisit the results based on FBA with respect to growth on myristate and export of Pi. Without additional constraints in the enzyme constrained iRi1574 model, the positive effect of myristate uptake on growth could not be reproduced, since myristate is catabolized via peroxisomal β-oxidation and the expression of the required enzymes is not outweighed by the benefit of generating acetyl-CoA from myristate. However, when the allocation of total protein is shifted from the optimal ratio towards increased abundances of peroxisomal proteins (suppl. Methods), the addition of myristate can increase growth compared to the suboptimal scenario (Figure S4). Further, we found that using the default medium, like in the FBA model above, the enzyme-constrained model predicts export of Pi at optimal growth in the range [0, 171.9] *mmol gDW*^−1^*h*^−1^ (Suppl. Tab 4). Therefore, the observations made for the FBA model with these important phenotypes also hold for the enzyme-constrained model.

To test the performance of the enzyme-constrained variant of the iRi1574 model, we made use of published dry weight and protein content available for 12 combinations of four carbon sources (i.e. D-glucose, D-fructose, raffinose, and melibiose) at three different concentrations (i.e. 10 mM, 100 mM, 1 M) (60). These data were generated by using the *G. intraradices* strain Sy167 (60), which is the closest species to *R. irregularis* for which this kind of measurements are available to date. The different media conditions were modelled by adding each carbohydrate to the default medium as a single carbon source, while the respective concentrations were modelled as proportional uptake fluxes considering kinetic parameters of the respective transport reactions (for more detail see Methods section). Like in the findings based on FBA, above, palmitate was present in the default medium since growth without palmitate is not possible, irrespective of additional supply of carbohydrates (45, 47). To avoid compensation of lower carbohydrate supply by β-oxidation of palmitic acid, we limited its uptake to the flux value obtained at optimal growth predicted by FBA.

We next compared the predictions of growth from the eMOMENT approach with those from FBA (i.e. without considering enzyme constraints), with the same restrictions on palmitate uptake (Suppl. Tab 5). We observed that the additional constraints on protein abundances largely improved the quality of the prediction (Fig. 2) and resulted in values of the same order as growth rates calculated from dry weights and grow duration (suppl. Methods). We found that the predicted growth rates were significantly correlated with the measured values for hyphae dry weight (Spearman correlation coefficient, *ρ_S_* = 0.74, *P* < 0.01, Fig. 2) and were collinear (*ρ_S_* = 1.0) with the protein content. In contrast, FBA predicted a statistically significant, negative correlation (*ρ_S_* = −0.69, *P* < 0.05) demonstrating that the predictions from this approach are not in line with the experimental observations. The respective values for Pearson correlation were *ρ_P_* = 0.80 (*P* < 0.01), for the enzyme-constrained models, and *ρ_P_* = −0.62 (*P* < 0.05), for the FBA model. Using FBA, we observed that growth increased with the concentration of the respective carbon source despite rescaling of biomass coefficients, while this was not the case when using the eMOMENT approach. In fact, this relationship was only observed for D-glucose and raffinose, which is broken down to sucrose and D-galactose extracellularly. Hence, the iRi1574 model can reliably predict growth based on different carbon sources when protein content and protein-reaction associations are considered.

**Figure 2.**
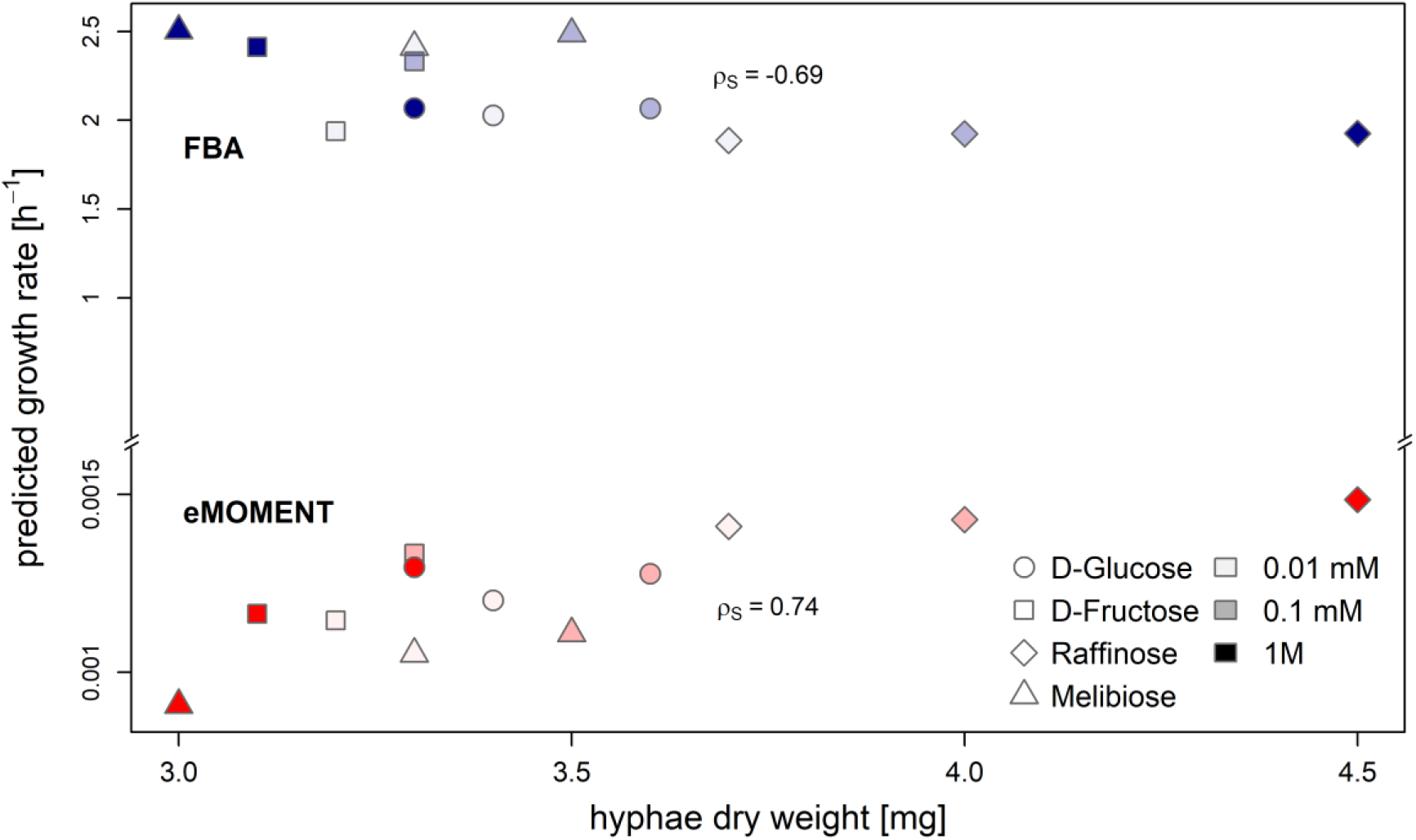
Prediction of growth for iRi1574 using eMOMENT and FBA. Scatter plot of growth rates predicted by eMOMENT (red) compared with FBA without constraints on enzyme abundances (blue). The predicted growth rates were compared with experimental data obtained for *Glomus intraradices* Sy167 (60), which is the phylogenetically closest species with this kind of data available. The concordance of predicted growth rates and experimentally measured hyphae dry weight was quantified by the Spearman correlation *ρ_S_*.

The applied approach to integrate total protein content allowed us to predict not only optimal growth, but also abundances of individual proteins for the 12 combinations of carbon source and concentrations considered. Since multiple allocations of proteins to enzyme complexes and reactions can lead to optimal growth, we sampled the set of feasible enzyme abundances (Methods) at 99% of the respective optima. The resulting predictions on alternative enzyme allocation at optimal growth were used to investigate the plasticity of enzyme allocation under the different conditions. We quantified the plasticity in the abundance of each protein by the coefficient of variation (CV) across the simulated conditions. The CV was calculated for predicted protein abundance and reaction flux across the 12 growth scenarios (Suppl. Tab 6 and 7). To illustrate the findings, we represented the distribution of CVs across the 13 metabolic subsystems (Fig. 3A-B). The highest median CV was found for enzymes within the amino sugar and nucleotide sugar metabolism 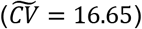, carbohydrate metabolism 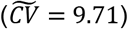, and nucleotide metabolism 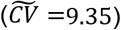. In contrast, metabolism of cofactors and vitamins and transport reactions showed the lowest plasticity in protein abundance 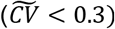. Regarding reaction fluxes, the subsystems containing the most plastic reactions were found within sink reactions 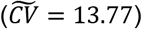 and amino sugar and nucleotide sugar metabolism 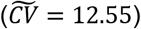.

**Figure 3.**
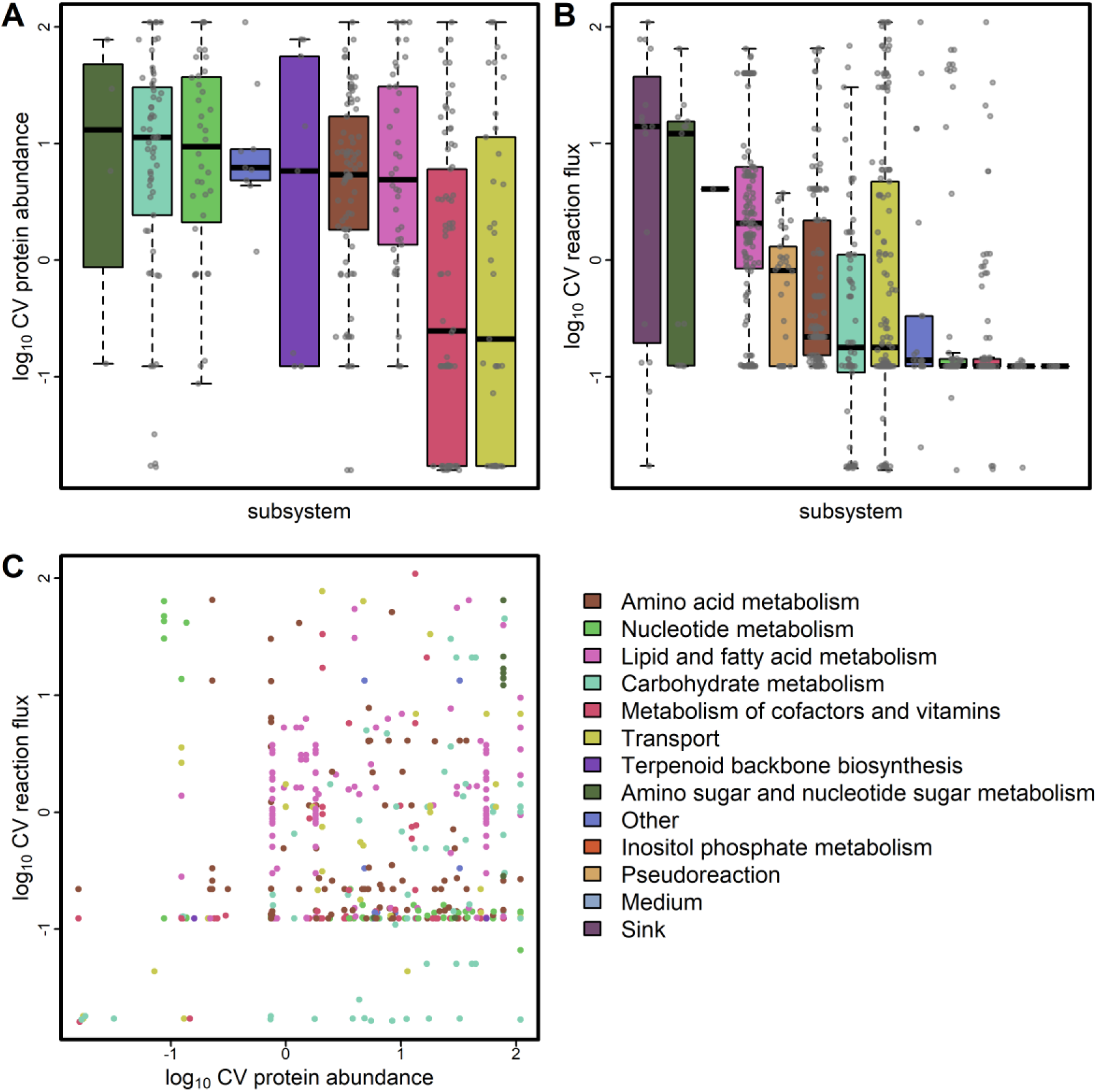
Plasticity of protein abundance and reaction fluxes across 12 simulated media conditions. The coefficient of variation (CV) was calculated across all media conditions (i.e. glucose, fructose, raffinose, and melibiose at 10, 100, and 1000 mM each) for protein abundances (**A**) and reaction fluxes (**B**). The boxes are ordered by median of the log10-transformed data. (**C**) The CV of fluxes is plotted against the CV of abundance of their associated proteins.

To assess whether the plasticity in flux is dependent on the variability in enzyme abundance of the catalysing enzymes, we compared the respective sets at the extreme ends of CV distribution (10% and 90% quantiles) between protein abundance and reaction fluxes (Figure 3C). Among the 89 reactions associated to enzymes with variable abundance (*CV* ≥ 49.38), 28 were found to be also highly plastic in flux. Conversely, we found six reactions with low flux CV associated with seven high abundance CV proteins. Four out of these genes were not promiscuous, and were associated with single reactions of high CV. The associated reactions are involved in terpenoid backbone biosynthesis and nucleotide metabolism. Hence, variation in enzyme abundance cannot fully explain the plasticity in flux. Since pH differences (affecting enzyme activity) are expected to lead to systemic changes, we conclude that the plasticity in flux for these selected reactions is largely driven by metabolite concentration, rather than enzyme abundance.

Among the 78 reactions with highly variable fluxes (*CV* ≥ 39.79), the majority lies within the lipid and fatty acid metabolism (47) and transport reactions (10). The subset of reactions in lipid metabolism were found to act mainly in in lipid degradation but also in the synthesis of very long chain fatty acids. This result indicates a trade-off between lipid synthesis and *β*-oxidation depending on the type and concentration of the carbon source.

### Prediction of growth for three fungal structures

As obligate biotrophs, AMF are dependent on the association with a host plant for carbohydrates and lipids (2, 3). Three major fungal structures are discriminated for the fungus: extraradical mycelium (ERM), intraradical mycelium (IRM) and arbuscles (ARB), which differ from each other in the proximity of association with the host plant. Thus, we investigated growth and underlying flux distributions comparing these three structures of *R. irregularis*. To this end, we used published expression data from (5) to examine growth and differential reaction fluxes between these three structures.

We observed an increase in growth upon association of the fungus with the host plant (Fig. 4A), which was expected since a tighter association with the host plant and hence increased nutrient uptake allows faster growth. Since the total protein content remained the same over the simulations for all three structures, the changes in growth likely result from increased flux through a subset of reactions responsible for growth, due to larger upper bounds of these reactions. One reason for this could be changes in the relative abundances of individual proteins due to changes in transcript abundances that were used to calculate the upper bounds. To determine differential reactions, we sampled 5000 flux distributions for each structure and compared the resulting flux values for each reaction using the non-parametric common-language effect size (*A_w_* (89), Suppl. Tab 8). We used three different thresholds for *A_w_* (i.e. 0.6, 0.7, 0.8) to find differentially activated reactions between each pair of structures. By using 0.6 as a threshold, we found that mainly reactions of the amino acid metabolism exhibited differential fluxes between each pair of structures, followed by reactions in metabolism of cofactors and vitamins, carbohydrate, lipid, and nucleotide metabolism (Fig. 4B). Upon increasing the threshold to 0.7, we found only two reactions to be differentially activated between the ERM compared IRM and ARB, which were both involved in riboflavin biosynthesis (KEGG M00911) (Fig. 4B). Moreover, eight reactions from metabolism of cofactors and vitamins were differential between IRM and ARB. When the threshold was increased to 0.8, only one reaction was found to differ between IRM and ARB, namely the coproporphyrinogen:oxygen oxidoreductase (E.C. 1.3.3.3). These results suggest that substantial rerouting of fluxes within these pathways might occur upon establishing the fungal-plant interface. However, differences in predicted growth may not exclusively result from large changes in for few reactions. It is likely, that small changes in a number of other reactions also contribute to an increased growth rate.

**Figure 4.**
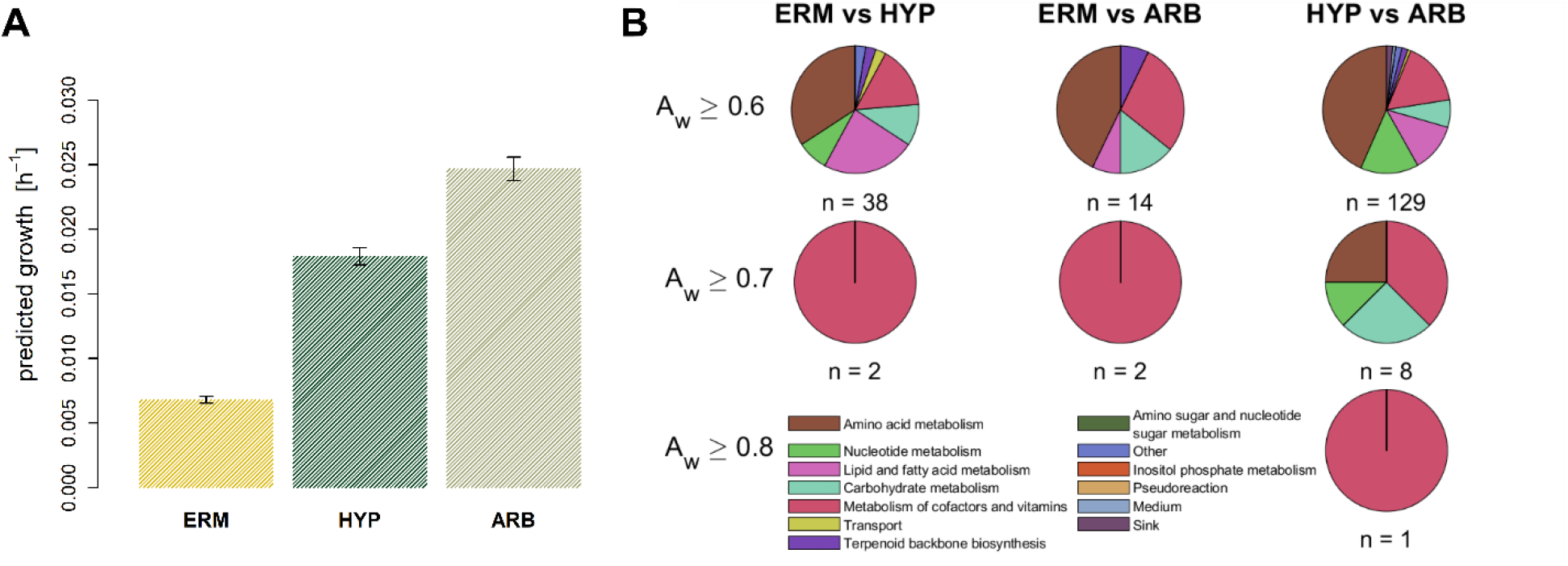
Growth simulation of *R. irregularis* for three fungal structures. The upper limit for reaction flux was calculated as *k_cat_* · [E]. Values for turnover constants associated to reaction was done similarly as in GECKO (68). Structure-specific expression data were used as proxy for protein concentrations. This was done by multiplying relative transcript abundances with the maximum total protein content measured with the available carbon source (*C* = 0.106 *g gDW*^−1^) (60) (see Methods section for more detail). **(A)** predicted growth for the three fungal structures. The error bars represent predicted growth rates at *C* ±*σ*, where *σ* represents the standard deviation determined for the experimentally measured protein content. **(B)** Distribution of subsystems for reactions that show non-parametric common language effect sizes (*A_w_*) above selected thresholds for each pairwise comparison of flux distributions between the three fungal structures. The total numbers of reactions with *A_w_* greater than the threshold are shown below each of the pie charts. No chart is shown if no reaction was found to have an *A_w_* above the threshold.

## Conclusion

Although *R. irregularis* is one of the most extensively studied AMF that forms symbioses with major crops, insights from the annotation of its enzymatic genes, the extensive body of evidence about its physiological and molecular responses to different environmental stimuli, and mutual effects on plants with which it interacts have not yet been systematically investigated in the context of metabolic modelling. The constraint-based modelling framework allows us to dissect the molecular mechanisms that underpin these responses and also to suggest targets for future metabolic engineering in order to boost the beneficial effects of this AMF. However, achieving this aim requires the assembly of a high-quality large-scale model that leads to accurate quantitative predictions of multiple traits in different scenarios.

Here, we presented the enzyme-constrained iRi1574 GEM of *R. irregularis* based on the KBase fungal reconstruction pipeline followed by consideration and inclusion of exhaustive literature research as well as manual curation for consistency, mass- and charge balance. One possible caveat of using fungal reconstruction pipelines is that the resulting model may be very similar to the employed templates. Nevertheless, by conducting comparative analysed of the enzyme set of iRi1574 and that of published fungal models, we demonstrated the specificity of iRi1574 and its ability to capture the particularities of *R. irregularis* metabolism. More importantly, validation tests demonstrated that iRi1574 can: (1) accurately predict increased growth on myristate with minimal medium with the FBA model as well as, under additional constrains on enzyme distribution, in the enzyme-constrained model, (2) predict growth that is highly correlated with hyphal dry weight measured in a close relative (*Glomus intraradices* Sy167, neighbouring clade), when considering enzyme constraints, and (3) growth rate increases with tighter association with the host plant, based on integration of relative transcriptomics data. The extensively validated model was used to show that the transition from IRM to ARB could be linked with changes in amino acid and cofactor biosynthesis.

This first model of an AMF can be coupled with root-specific models of model plants to investigate the effects of symbiosis. Further, a 2D growth simulation approach (62) can be employed to obtain a realistic growth measure for hyphal spread. In addition, the iRi1574 model can be used to mechanistically dissect the interactions of species in fungal and bacterial communities that jointly affect plant performance (90). Most importantly, one can begin to design metabolic engineering strategies to improve desired traits in *R. irregularis*, study the effect of the modifications on plant performance by coupling metabolic models of the symbionts, and to further refine the model based on integration of heterogeneous molecular data. Altogether these modelling efforts can guide future reverse genetics tools used to understand the functional relevance of metabolic genes in *R. irregularis* in shaping plant traits.

## Materials and Methods

### Draft model generation

The genome of *Rhizophagus irregularis* DAOM 181602=DAOM 197198 (GCF_000439145.3) (49, 51) served as the basis for the genome-scale metabolic reconstruction. The initial draft model was obtained from KBase (63) using the ‘Build Fungal Model’ app (Oct 15, 2018; narrative ID 36938). The resulting model was gap-filled with the help of the KBase app ‘Gapfill Metabolic Model’ app using the complete medium. A set of 35 additional reactions was required to simulate growth. This set of added reactions was manually curated in the next step of model refinement. The gap-filled model was then downloaded in SBML format and further modified within MATLAB (91) using functions of the COBRA toolbox (92).

### Model curation

To enhance connectivity between the cellular compartments, 198 transport reactions were added from the yeast iMM904 metabolic model (64). The imported transport reactions were validated during the next curation steps. Out of all added transport reactions, 71 were kept in the model despite missing literature evidence (Suppl. Tab 9). Next, the metabolite and reaction identifiers were translated, whenever possible, to the ModelSEED namespace (34). This step was necessary since the identifiers resulted from 14 different models and the catalyzed reactions mostly could not be identified. Moreover, this led to a higher connectivity of the network as identical metabolites and reactions were reconciled. Further, cross-references were added to BiGG (93), MetaCyc (94), KEGG (74), MetaNetX (95), PubChem (96), and E.C. numbers.

Metabolite formulas were added from PubChem and adapted to the net charge at the average cytosolic pH of 6.2 (97) using the ChemAxon Marvin software (Marvin 17.21.0, 2017, http://www.chemaxon.com). With elemental compositions and metabolite charges available, the model was manually mass- and charge balanced.

After these steps, additional reactions were added from various literature sources. Most of the lipid metabolism is based on the results from (45), including: Sterol metabolism, Fatty acid synthesis, -elongation, and -degradation, glycerolipid metabolism, sphingolipid metabolism. Plasma membrane transporters were added with literature evidence from multiple sources (38, 44, 53, 98). Furthermore, important dead-end metabolites were resolved manually by adding incident reactions with genetic evidence or transport reactions.

The biomass reaction was adapted from the default fungal biomass reaction added during the automated reconstruction process (Suppl. Tab 10). Subsequently, the unknown coefficients in the biomass reaction were re-scaled such that the sum of coefficients multiplied with the respective molecular weight equals 1 *g gDW*^−1^ (99). Due to missing experimental data, we set the growth-associated ATP maintenance reaction (GAM) to 60 molecules ATP *gDW*^−1^ as taken from the KBase default fungal biomass, which is in line with the average value from seven published fungal models (68.87 *mmol gDW*^−1^*h*^−1^, Suppl. Tab 11). The non-growth associated ATP maintenance reaction (NGAM) was fixed to the average of from seven published fungal models (3.21 *mmol gDW*^−1^*h*^−1^, Suppl. Tab 11). For the lipid component in the biomass reactions, the SLIMEr formalism was used (100) and coefficients of tail and backbone pseudometabolites were adjusted to render the model feasible for simulations by running a quadratic program to minimize factors to be added to the respective coefficients.

Stoichiometrically-balanced cycles (SBC) were then removed by repeatedly applying Flux Variability Analysis (FVA) and correcting reaction reversibility or adding additional reactions as suggested in (101). For the following analyses, all reversible reactions were split into two irreversible reactions.

### Short-chain chitooligosaccharides (CO) and lipochitooligosaccharides (LCO)

Synthesis reactions for LCOs were added by first modelling the synthesis of COs with chain lengths 3-6 with subsequent acetylation reactions adding 16:0, 16:1*Δ*9(*ω*7), 18:0, and 18:1*Δ*9(*ω*9) fatty acids leading to 16 different LCO species (70, 71).

### Transcriptomic data

Structure-dependent RNA-seq data were obtained as raw sequence reads (GSE99655) (5). The reads were quality trimmed using Trimmomatic-0.39 (102) and mapped to the *R. irregularis* genome using STAR 2.7.3a (103). The read quantification was performed using HTSeq count (104). The average over the three replicates was used for further analysis. The protein identifiers from the original study were translated to the identifiers of the genome annotation that was used for the metabolic reconstruction using local tblastn (105, 106) using the BLOSUM90 scoring matrix and a cutoff E-value of 10E-90. The average Spearman correlation between the published and re-analysed values for the secreted proteins (SP) was *0.8*, which confirms the previous results given different analysis software and possible mapping errors using tblastn.

### Turnover numbers

For the assignment of *k_cat_* values to reactions, a similar approach as in GECKO (68) was applied. First, turnover values for all E.C. numbers in the model across all organisms and according lineages were obtained from BRENDA (107), SABIO-RK (108) and UniProt (109), respectively. For each E.C. number assigned to a reaction, all matching *k_cat_* values were obtained and, if possible, filtered for substrate matches and enzymes from the fungi kingdom. If no match for the complete E.C. number was found the same procedure was applied to the same E.C. number pruned to a lower level. Among the obtained values, the maximum *k_cat_* value was assigned to the respective reaction. The distribution and numbers of matched *k_cat_* values per subsystem, as well as a comparison to *k_cat_* values in the YestGEM v8.3.4 are shown in Figures S5A and B. The median of all non-zero values was used for metabolic reactions without a matched *k_cat_* value. To arrive at units of *h*^−1^, all turnover numbers were multiplied by 3600.

### Enzyme usage under different growth conditions

To predict the enzyme abundances with different media conditions, four different carbon sources (i.e. D-glucose, D-fructose, raffinose, melibiose) were added to the minimal medium (65) (Suppl. Tab 12) as single carbon sources. These carbohydrates were selected, as hyphal weight and protein content were available for them at three different concentrations (i.e. 10, 100, and 1000 *mM*) (60). As an exception, palmitate was retained in the medium as it must be supplied to the fungus in order to allow for growth (45, 47). We used kinetic parameters (i.e. *V_max_* and *K_m_*) of *S. cerevisiae* monosaccharide transporters to model the influx of D-glucose, D-fructose, and D-galactose (results from breakdown of both raffinose and melibiose, Suppl. Tab 13) (110, 111). The respective upper bound for the transporters was calculated as

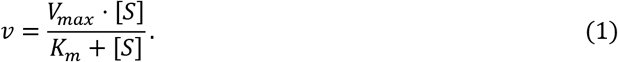

Further, the import of palmitate was restricted to the flux value of 8.46 *mmol gDW*^−1^*h*^−1^ at optimal growth as predicted by FBA.

The following MILP, which we termed eMOMENT, imposes constraints which were adopted from the MOMENT approach (84), which were extended by an additional constraint (Eq. 4):

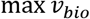

**s.t.**

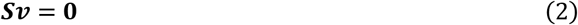

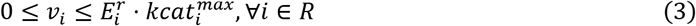

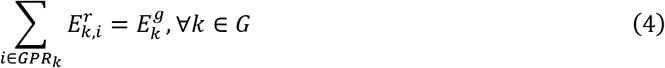

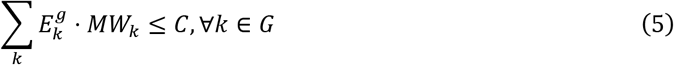

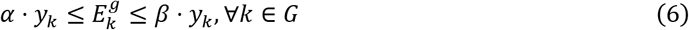

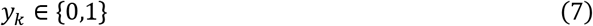

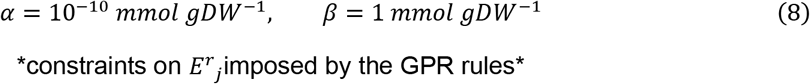

*R* and *G* represent the sets of reactions and genes. The molecular weight in *g mmol*^−1^ of a protein *k* is given by *MW_k_*. The constraint in Eq. (3) imposes an upper limit on the flux through reaction *i* which is the product of the reaction-specific turnover rate and the enzyme abundance *E^r^_i_* available for this reaction. Further, binary variables *y* were introduced to indicate that the respective genes are expressed (*y* = 1) or not (*y* = 0). This was done to enforce a lower bound *α* for the abundance of expressed genes to avoid numerical problems. The value for *E^r^_i_* is determined by the GPR rules. To model the GPR rules, the following constraints were applied recursively in case of complex rules:

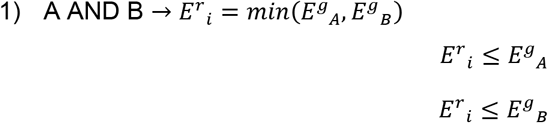

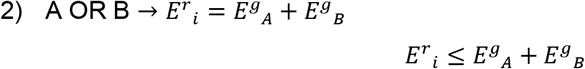

Further, the total protein content *C* was determined by the experimentally-measured protein contents at the given concentrations (60). To account for changing protein contents, the coefficients of the biomass reaction were rescaled to the respective values for *C*. The proportions of the remaining biomass components were conserved when they were adapted to the new residual mass fraction (1 *g gDW*^−1^ – *C*).

We extended the constraints we borrowed from the MOMENT approach by one additional constraint (Eq. (4)), which takes the promiscuity of proteins for multiple reactions into account. Hence, the abundance of protein *k* is smaller than or equal to the sum of enzyme abundances across all reactions with which it is associated.

The feasible abundance ranges for all proteins were determined by individual minimization and maximization for *E^g^_i_*, at optimal growth, similar to FVA. Using these, we sampled 1000 abundances compatible with the constraints above, by finding the closest vector of abundances to a randomly created set of abundances *E^g^*\* within the feasible ranges determined in the step before:

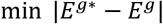

**s.t.**

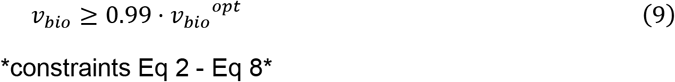

### Metabolic changes between fungal structures

For this experiment, all four carbon sources that were used in the analysis above, were added to the same minimal medium. Similarly, the upper bounds on monosaccharide import were calculated using transporter kinetics from *S. cerevisiae*, considering only the maximum concentration of *1 M*. Across the calculated values, the maximum possible influx for each monosaccharide was selected. For this experiment, palmitate was also retained in the medium with the same upper limit as described before. For each of the three structures (ERM, IRM, ARB), the abundance of each protein *tc^p^_i_* was calculated from the relative transcriptomic counts per gene *tc^g^_i_* (not considering alternative splicing and post-translational modifications):

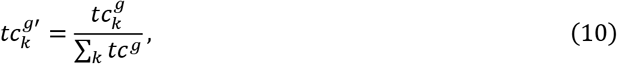

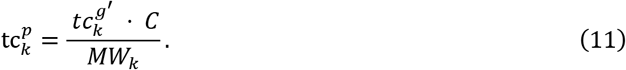

The total protein content *C* was set to the maximum value measured across all growth conditions used in the experiment before (*C* = 0.106 *g gDW*^−1^). By applying this transformation, we assume that transcript levels correlate with protein abundances, which is not necessarily true and can lead to over- or underestimation of protein levels. However, this represents the closest approximation of protein levels in the absence of quantitative proteomics data.

To conduct FBA, the transformed transcript count *tc^r^_i_* for the reaction was first calculated by applying the GPR rules taking the minimum *tc^p^* value for complexes (AND) and the maximum for isozymes (OR). Finally, the upper limit for a reaction *i* was defined as the product of estimated enzyme abundance and the respective turnover value:

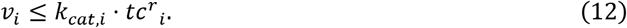

Growth was predicted for each of the three structures by FBA using the adapted reaction limits. After this, FVA was used to determine the feasible ranges for each reaction while keeping the growth at 99% of the optimum. These ranges were used as the limits for the sampling procedure which attempts to find an optimal solution with minimal distance to a random flux vector *ν*\*:

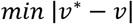

**s.t.**

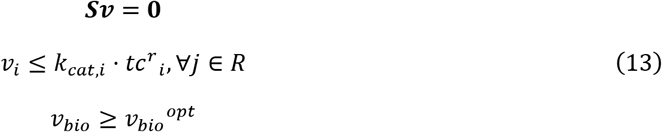

Like this, 5000 points were sampled and used for a reaction-wise comparison between the three structures. To this end, the non-parametric estimate for common language *A_w_* (89) was used to determine substantial changes of reaction flux between each pair of structures:

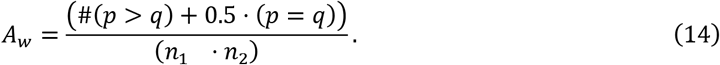

The variables *p* and *q* represent the vectors of sampled fluxes for the same reaction at two different structures.

## Data and code availability

All procedures, data, and approaches used are available at https://github.com/pwendering/RhiirGEM.

## Acknowledgments

Z.N would like to acknowledge the support from the Max Planck Society. Z.N. and P.W. would like to acknowledge the funding from the Research Focus “Evolutionary Systems Biology” of the University of Potsdam.

## Supplementary Information Legends

**Supplementary File S1:** iRi1574 model in Systems Biology Markup Language (SBML) format.

**Supplementary File S2:** Quality assessment report from the MEMOTE test suite (76).

**Supplementary Table S1:** iRi1574 model in Excel format.

**Supplementary Table S2:** Characteristics of published fungal models selected for comparison with iRi1574.

**Supplementary Table S3:** FBA solution and feasible ranges and of all reactions in the iRi1574 model determined by FVA.

**Supplementary Table S4:** eMOMENT solution and feasible ranges and of all reactions in the enzyme-constraint iRi1574 model.

**Supplementary Table S5:** Experimentally measured hyphal dry weight and protein content as well as growth predictions by FBA and eMOMENT with different saccharides and their concentrations (60).

**Supplementary Table S6:** Coefficients of variation (CV) of abundances for each protein across all simulated conditions (carbon source x concentration).

**Supplementary Table S7:** Coefficients of variation (CV) of fluxes for each reaction across all simulated conditions (carbon source x concentration).

**Supplementary Table S8:** Non-parametric common language effect sizes between fluxes at different developmental stages.

**Supplementary Table S9:** Transport reactions in the iRi1574 model that were introduced from the iMM904 model and retained without literature evidence.

**Supplementary Table S10:** Biomass composition of the iRi1574 model.

**Supplementary Table S11:** Flux limits for the (non-)growth associated ATP maintenance reactions (NGAM, GAM) from fungal models.

**Supplementary Table S12:** Minimal medium used for simulations.

**Supplementary Table S13:** Calculation of hexose influxes for each of the carbon sources that were used in the 12 simulated scenarios.

**Supplementary Table S14:** Influence of reported SBC reactions on predicted growth.

## References

1. Smith S, Read D. 2008. Mycorrhizal Symbiosis, 3rd ed. Academic Press, London.

2. Parniske M. 2008. Arbuscular mycorrhiza: The mother of plant root endosymbioses. Nat Rev Microbiol. Nat Rev Microbiol.

3. Wipf D, Krajinski F, van Tuinen D, Recorbet G, Courty PE. 2019. Trading on the arbuscular mycorrhiza market: from arbuscules to common mycorrhizal networks. New Phytol 223:1127–1142.

4. Stockinger H, Walker C, Schüßler A. 2009. “*Glomus intraradices* DAOM197198”, a model fungus in arbuscular mycorrhiza research, is not *Glomus intraradices*. New Phytol 183:1176–1187.

5. Zeng T, Holmer R, Hontelez J, Lintel-Hekkert B, Marufu L, Zeeuw T, Wu F, Schijlen E, Bisseling T, Limpens E. 2018. Host- and stage-dependent secretome of the arbuscular mycorrhizal fungus *Rhizophagus irregularis*. Plant J 94:411–425.

6. Zhao S, Chen A, Chen C, Li C, Xia R, Wang X. 2019. Transcriptomic analysis reveals the possible roles of sugar metabolism and export for positive mycorrhizal growth responses in soybean. Physiol Plant 166:712–728.

7. Li Z, Ngwene B, Hong T, George E. 2019. Effects of nitrogen feeding for extraradical mycelium of *Rhizophagus irregularis* maize symbiosis incorporated with phosphorus availability. J Plant Nutr Soil Sci 182:647–655.

8. Yang Q, Ravnskov S, Neumann Andersen M. 2020. Nutrient uptake and growth of potato: Arbuscular mycorrhiza symbiosis interacts with quality and quantity of amended biochars. J Plant Nutr Soil Sci 183:220–232.

9. Chaudhary V, Kapoor R, Bhatnagar AK. 2008. Effectiveness of two arbuscular mycorrhizal fungi on concentrations of essential oil and artemisinin in three accessions of *Artemisia annua* L. Appl Soil Ecol 40:174–181.

10. Tekaya M, Mechri B, Mbarki N, Cheheb H, Hammami M, Attia F. 2017. Arbuscular mycorrhizal fungus Rhizophagus irregularis influences key physiological parameters of olive trees (*Olea europaea* L.) and mineral nutrient profile. Photosynthetica 55:308–316.

11. Wahbi S, Prin Y, Maghraoui T, Sanguin H, Thioulouse J, Oufdou K, Hafidi M, Duponnois R. 2015. Field Application of the Mycorrhizal Fungus *Rhizophagus irregularis* Increases the Yield of Wheat Crop and Affects Soil Microbial Functionalities. Am J Plant Sci 06:3205–3215.

12. Goicoechea N, Bettoni MM, Fuertes-Mendizábal T, González-Murua C, Aranjuelo I. 2016. Durum wheat quality traits affected by mycorrhizal inoculation, water availability and atmospheric CO2 concentration. Crop Pasture Sci 67:147.

13. Todeschini V, Aitlahmidi N, Mazzucco E, Marsano F, Gosetti F, Robotti E, Bona E, Massa N, Bonneau L, Marengo E, Wipf D, Berta G, Lingua G. 2018. Impact of beneficial microorganisms on strawberry growth, fruit production, nutritional quality, and volatilome. Front Plant Sci 9:1611.

14. Navarro JM, Morte A. 2019. Mycorrhizal effectiveness in *Citrus macrophylla* at low phosphorus fertilization. J Plant Physiol 232:301–310.

15. Rivera-Becerril F. 2002. Cadmium accumulation and buffering of cadmium-induced stress by arbuscular mycorrhiza in three *Pisum sativum* L. genotypes. J Exp Bot 53:1177–1185.

16. Porras-Soriano A, Soriano-Martín ML, Porras-Piedra A, Azcón R. 2009. Arbuscular mycorrhizal fungi increased growth, nutrient uptake and tolerance to salinity in olive trees under nursery conditions. J Plant Physiol 166:1350–1359.

17. Cattani I, Beone GM, Gonnelli C. 2015. Influence of *Rhizophagus irregularis* inoculation and phosphorus application on growth and arsenic accumulation in maize (*Zea mays* L.) cultivated on an arsenic-contaminated soil. Environ Sci Pollut Res 22:6570–6577.

18. Zhu X, Song F, Liu F. 2016. Altered amino acid profile of arbuscular mycorrhizal maize plants under low temperature stress. J Plant Nutr Soil Sci 179:186–189.

19. Calvo-Polanco M, Sánchez-Romera B, Aroca R, Asins MJ, Declerck S, Dodd IC, Martínez-Andújar C, Albacete A, Ruiz-Lozano JM. 2016. Exploring the use of recombinant inbred lines in combination with beneficial microbial inoculants (AM fungus and PGPR) to improve drought stress tolerance in tomato. Environ Exp Bot 131:47–57.

20. Zhang H, Wei S, Hu W, Xiao L, Tang M. 2017. Arbuscular Mycorrhizal Fungus *Rhizophagus irregularis* Increased Potassium Content and Expression of Genes Encoding Potassium Channels in *Lycium barbarum*. Front Plant Sci 8:1–11.

21. Garg N, Singh S. 2018. Arbuscular Mycorrhiza *Rhizophagus irregularis* and Silicon Modulate Growth, Proline Biosynthesis and Yield in *Cajanus cajan* L. Millsp. (pigeonpea) Genotypes Under Cadmium and Zinc Stress. J Plant Growth Regul 37:46–63.

22. Begum N, Qin C, Ahanger MA, Raza S, Khan MI, Ashraf M, Ahmed N, Zhang L. 2019. Role of Arbuscular Mycorrhizal Fungi in Plant Growth Regulation: Implications in Abiotic Stress Tolerance. Front Plant Sci 10.

23. Smith SE, Smith FA, Jakobsen I. 2003. Mycorrhizal Fungi Can Dominate Phosphate Supply to Plants Irrespective of Growth Responses. Plant Physiol 133:16–20.

24. Govindarajulu M, Pfeffer PE, Jin H, Abubaker J, Douds DD, Allen JW, Bücking H, Lammers PJ, Shachar-Hill Y. 2005. Nitrogen transfer in the arbuscular mycorrhizal symbiosis. Nature 435:819–823.

25. Cruz C, Egsgaard H, Trujillo C, Ambus P, Requena N, Martins-Loução MA, Jakobsen I. 2007. Enzymatic evidence for the key role of arginine in nitrogen translocation by arbuscular mycorrhizal fungi. Plant Physiol 144:782–792.

26. Fiorilli V, Lanfranco L, Bonfante P. 2013. The expression of GintPT, the phosphate transporter of *Rhizophagus irregularis*, depends on the symbiotic status and phosphate availability. Planta 237:1267–1277.

27. Ezawa T, Saito K. 2018. How do arbuscular mycorrhizal fungi handle phosphate? New insight into fine-tuning of phosphate metabolism. New Phytol. Blackwell Publishing Ltd.

28. Ho I, Trappe JM. 1973. Translocation of ^14^C from *Festuca* Plants to their Endomycorrhizal Fungi. Nat New Biol 244:30–31.

29. Solaiman MZ, Saito M. 1997. Use of sugars by intraradical hyphae of arbuscular mycorrhizal fungi revealed by radiorespirometry. New Phytol 136:533–538.

30. Bago B, Pfeffer PE, Shachar-Hill Y. 2000. Carbon Metabolism and Transport in Arbuscular Mycorrhizas. Plant Physiol 124:949–958.

31. Ait Lahmidi N, Courty PE, Brulé D, Chatagnier O, Arnould C, Doidy J, Berta G, Lingua G, Wipf D, Bonneau L. 2016. Sugar exchanges in arbuscular mycorrhiza: RiMST5 and RiMST6, two novel *Rhizophagus irregularis* monosaccharide transporters, are involved in both sugar uptake from the soil and from the plant partner. Plant Physiol Biochem 107:354–363.

32. Jiang Y, Wang W, Xie Q, Liu N, Liu L, Wang D, Zhang X, Yang C, Chen X, Tang D, Wang E. 2017. Plants transfer lipids to sustain colonization by mutualistic mycorrhizal and parasitic fungi. Science (80-) 356:1172–1173.

33. Luginbuehl LH, Menard GN, Kurup S, Van Erp H, Radhakrishnan G V., Breakspear A, Oldroyd GED, Eastmond PJ. 2017. Fatty acids in arbuscular mycorrhizal fungi are synthesized by the host plant. Science (80-) 356:1175–1178.

34. Henry CS, DeJongh M, Best AA, Frybarger PM, Linsay B, Stevens RL. 2010. High-throughput generation, optimization and analysis of genome-scale metabolic models. Nat Biotechnol 28:977–982.

35. Keymer A, Pimprikar P, Wewer V, Huber C, Brands M, Bucerius SL, Delaux PM, Klingl V, von Röpenack-Lahaye E, Wang TL, Eisenreich W, Dörmann P, Parniske M, Gutjahr C. 2017. Lipid transfer from plants to arbuscular mycorrhiza fungi. Elife 6:1–33.

36. Koide RT, Kabir Z. 2000. Extraradical hyphae of the mycorrhizal fungus *Glomus intraradices* can hydrolyse organic phosphate. New Phytol 148:511–517.

37. Maldonado-Mendoza IE, Dewbre GR, Harrison MJ. 2001. A phosphate transporter gene from the extra-radical mycelium of an arbuscular mycorrhizal fungus *Glomus intraradices* is regulated in response to phosphate in the environment. Mol Plant-Microbe Interact 14:1140–1148.

38. Walder F, Boller T, Wiemken A, Courty PE. 2016. Regulation of plants’ phosphate uptake in common mycorrhizal networks: Role of intraradical fungal phosphate transporters. Plant Signal Behav 11:e1131372.

39. Courty PE, Smith P, Koegel S, Redecker D, Wipf D. 2015. Inorganic Nitrogen Uptake and Transport in Beneficial Plant Root-Microbe Interactions. CRC Crit Rev Plant Sci 34:4–16.

40. López-Pedrosa A, González-Guerrero M, Valderas A, Azcón-Aguilar C, Ferrol N. 2006. GintAMT1 encodes a functional high-affinity ammonium transporter that is expressed in the extraradical mycelium of *Glomus intraradices*. Fungal Genet Biol 43:102–110.

41. Pérez-Tienda J, Testillano PS, Balestrini R, Fiorilli V, Azcón-Aguilar C, Ferrol N. 2011. GintAMT2, a new member of the ammonium transporter family in the arbuscular mycorrhizal fungus *Glomus intraradices*. Fungal Genet Biol 48:1044–1055.

42. Calabrese S, Pérez-Tienda J, Ellerbeck M, Arnould C, Chatagnier O, Boller T, Schüßler A, Brachmann A, Wipf D, Ferrol N, Courty P-E. 2016. GintAMT3 – a Low-Affinity Ammonium Transporter of the Arbuscular Mycorrhizal *Rhizophagus irregularis*. Front Plant Sci 7:679.

43. Tian C, Kasiborski B, Koul R, Lammers PJ, Bucking H, Shachar-Hill Y. 2010. Regulation of the nitrogen transfer pathway in the arbuscular mycorrhizal symbiosis: Gene characterization and the coordination of expression with nitrogen flux. Plant Physiol 153:1175–1187.

44. Helber N, Wippel K, Sauer N, Schaarschmidt S, Hause B, Requena N. 2011. A versatile monosaccharide transporter that operates in the arbuscular mycorrhizal fungus *Glomus* sp is crucial for the symbiotic relationship with plants. Plant Cell 23:3812–3823.

45. Wewer V, Brands M, Dörmann P. 2014. Fatty acid synthesis and lipid metabolism in the obligate biotrophic fungus *Rhizophagus irregularis* during mycorrhization of *Lotus japonicus*. Plant J 79:398–412.

46. Roth R, Paszkowski U. 2017. Plant carbon nourishment of arbuscular mycorrhizal fungi. Curr Opin Plant Biol.

47. Sugiura Y, Akiyama R, Tanaka S, Yano K, Kameoka H, Marui S, Saito M, Kawaguchi M, Akiyama K, Saito K. 2020. Myristate can be used as a carbon and energy source for the asymbiotic growth of arbuscular mycorrhizal fungi. Proc Natl Acad Sci 117:202006948.

48. Abdellatif L, Lokuruge P, Hamel C. 2019. Axenic growth of the arbuscular mycorrhizal fungus *Rhizophagus irregularis* and growth stimulation by coculture with plant growth-promoting rhizobacteria. Mycorrhiza 29:591–598.

49. Tisserant E, Malbreil M, Kuo A, Kohler A, Symeonidi A, Balestrini R, Charron P, Duensing N, Frei dit Frey N, Gianinazzi-Pearson V, Gilbert LB, Handa Y, Herr JR, Hijri M, Koul R, Kawaguchi M, Krajinski F, Lammers PJ, Masclaux FG, Murat C, Morin E, Ndikumana S, Pagni M, Petitpierre D, Requena N, Rosikiewicz P, Riley R, Saito K, San Clemente H, Shapiro H, van Tuinen D, Becard G, Bonfante P, Paszkowski U, Shachar-Hill YY, Tuskan GA, Young JPW, Sanders IR, Henrissat B, Rensing SA, Grigoriev I V., Corradi N, Roux C, Martin F. 2013. Genome of an arbuscular mycorrhizal fungus provides insight into the oldest plant symbiosis. Proc Natl Acad Sci 110:20117–20122.

50. Lin K, Limpens E, Zhang Z, Ivanov S, Saunders DGO, Mu D, Pang E, Cao H, Cha H, Lin T, Zhou Q, Shang Y, Li Y, Sharma T, van Velzen R, de Ruijter N, Aanen DK, Win J, Kamoun S, Bisseling T, Geurts R, Huang S. 2014. Single Nucleus Genome Sequencing Reveals High Similarity among Nuclei of an Endomycorrhizal Fungus. PLoS Genet 10.

51. Chen ECH, Morin E, Beaudet D, Noel J, Yildirir G, Ndikumana S, Charron P, St-Onge C, Giorgi J, Krüger M, Marton T, Ropars J, Grigoriev I V., Hainaut M, Henrissat B, Roux C, Martin F, Corradi N. 2018. High intraspecific genome diversity in the model arbuscular mycorrhizal symbiont *Rhizophagus irregularis*. New Phytol 220:1161–1171.

52. Morin E, Miyauchi S, San Clemente H, Chen ECH, Pelin A, Providencia I, Ndikumana S, Beaudet D, Hainaut M, Drula E, Kuo A, Tang N, Roy S, Viala J, Henrissat B, Grigoriev I V., Corradi N, Roux C, Martin FM. 2019. Comparative genomics of *Rhizophagus irregularis*, *R. cerebriforme*, *R. diaphanus* and *Gigaspora rosea* highlights specific genetic features in Glomeromycotina. New Phytol 222:1584–1598.

53. Tamayo E, Gómez-Gallego T, Azcón-Aguilar C, Ferrol N. 2014. Genome-wide analysis of copper, iron and zinc transporters in the arbuscular mycorrhizal fungus *Rhizophagus irregularis*. Front Plant Sci 5:1–13.

54. Handa Y, Nishide H, Takeda N, Suzuki Y, Kawaguchi M, Saito K. 2015. RNA-seq Transcriptional Profiling of an Arbuscular Mycorrhiza Provides Insights into Regulated and Coordinated Gene Expression in *Lotus japonicus* and *Rhizophagus irregularis*. Plant Cell Physiol 56:1490–1511.

55. Calabrese S, Kohler A, Niehl A, Veneault-Fourrey C, Boller T, Courty PE. 2017. Transcriptome analysis of the *Populus trichocarpa-Rhizophagus irregularis* mycorrhizal symbiosis: Regulation of plant and fungal transportomes under nitrogen starvation. Plant Cell Physiol 58:1003–1017.

56. Calabrese S, Cusant L, Sarazin A, Niehl A, Erban A, Brulé D, Recorbet G, Wipf D, Roux C, Kopka J, Boller T, Courty PE. 2019. Imbalanced Regulation of Fungal Nutrient Transports According to Phosphate Availability in a Symbiocosm Formed by Poplar, Sorghum, and *Rhizophagus irregularis*. Front Plant Sci 10:1617.

57. Fang X, Lloyd CJ, Palsson BO. 2020. Reconstructing organisms in silico: genome-scale models and their emerging applications. Nat Rev Microbiol 18:1–13.

58. Pfau T, Christian N, Masakapalli SK, Swee LJ, Poolman MG, Ebenhöh O. 2018. The intertwined metabolism during symbiotic nitrogen fixation elucidated by metabolic modelling. Nature 8:1–11.

59. diCenco GC, Tesi M, Pfau T, Mengoni A, Fondi M. 2020. Genome-scale metabolic reconstruction of the symbiosis between a leguminous plant and a nitrogen-fixing bacterium. Nat Commun 11.

60. Hildebrandt U, Ouziad F, Marner F-J, Bothe H. 2006. The bacterium *Paenibacillus validus* stimulates growth of the arbuscular mycorrhizal fungus *Glomus intraradices* up to the formation of fertile spores. FEMS Microbiol Lett 254:258–267.

61. Bordbar A, Monk JM, King ZA, Palsson BO. 2014. Constraint-based models predict metabolic and associated cellular functions. Nat Rev Genet. Nature Publishing Group.

62. Harcombe WR, Riehl WJ, Dukovski I, Granger BR, Betts A, Lang AH, Bonilla G, Kar A, Leiby N, Mehta P, Marx CJ, Segrè D. 2014. Metabolic Resource Allocation in Individual Microbes Determines Ecosystem Interactions and Spatial Dynamics. Cell Rep 7:1104–1115.

63. Arkin AP, Cottingham RW, Henry CS, Harris NL, Stevens RL, Maslov S, Dehal P, Ware D, Perez F, Canon S, Sneddon MW, Henderson ML, Riehl WJ, Murphy-Olson D, Chan SY, Kamimura RT, Kumari S, Drake MM, Brettin TS, Glass EM, Chivian D, Gunter D, Weston DJ, Allen BH, Baumohl J, Best AA, Bowen B, Brenner SE, Bun CC, Chandonia JM, Chia JM, Colasanti R, Conrad N, Davis JJ, Davison BH, DeJongh M, Devoid S, Dietrich E, Dubchak I, Edirisinghe JN, Fang G, Faria JP, Frybarger PM, Gerlach W, Gerstein M, Greiner A, Gurtowski J, Haun HL, He F, Jain R, Joachimiak MP, Keegan KP, Kondo S, Kumar V, Land ML, Meyer F, Mills M, Novichkov PS, Oh T, Olsen GJ, Olson R, Parrello B, Pasternak S, Pearson E, Poon SS, Price GA, Ramakrishnan S, Ranjan P, Ronald PC, Schatz MC, Seaver SMDD, Shukla M, Sutormin RA, Syed MH, Thomason J, Tintle NL, Wang D, Xia F, Yoo H, Yoo S, Yu D. 2018. KBase: The United States Department of Energy Systems Biology Knowledgebase. Nat Biotechnol 36:566–569.

64. Mo ML, Palsson B, Herrgård MJ. 2009. Connecting extracellular metabolomic measurements to intracellular flux states in yeast. BMC Syst Biol 3.

65. Rosikiewicz P, Bonvin J, Sanders IR. 2017. Cost-efficient production of in vitro *Rhizophagus irregularis*. Mycorrhiza 27:477–486.

66. Vijayakumar V, Liebisch G, Buer B, Xue L, Gerlach N, Blau S, Schmitz J, Bucher M. 2016. Integrated multi-omics analysis supports role of lysophosphatidylcholine and related glycerophospholipids in the Lotus japonicus-Glomus intraradices mycorrhizal symbiosis. Plant Cell Environ 39:393–415.

67. Olsson PA, Johansen A. 2000. Lipid and fatty acid composition of hyphae and spores of arbuscular mycorrhizal fungi at different growth stages. Mycol Res 104:429–434.

68. Sánchez BJ, Zhang C, Nilsson A, Lahtvee P, Kerkhoven EJ, Nielsen J. 2017. Improving the phenotype predictions of a yeast genome-scale metabolic model by incorporating enzymatic constraints. Mol Syst Biol 13:935.

69. Pons S, Fournier S, Chervin C, Bécard G, Rochange S, Dit Frey NF, Pagès VP. 2020. Phytohormone production by the arbuscular mycorrhizal fungus Rhizophagus irregularis. PLoS One 15.

70. Maillet F, Poinsot V, André O, Puech-Pagés V, Haouy A, Gueunier M, Cromer L, Giraudet D, Formey D, Niebel A, Martinez EA, Driguez H, Bécard G, Dénarié J. 2011. Fungal lipochitooligosaccharide symbiotic signals in arbuscular mycorrhiza. Nature 469:58–64.

71. Genre A, Chabaud M, Balzergue C, Puech-Pagès V, Novero M, Rey T, Fournier J, Rochange S, Bécard G, Bonfante P, Barker DG. 2013. Short-chain chitin oligomers from arbuscular mycorrhizal fungi trigger nuclear Ca ^2+^ spiking in *Medicago truncatula* roots and their production is enhanced by strigolactone. New Phytol 198:190–202.

72. Callow JA, Capaccio LCM, Parish G, Tinker PB. 1978. Detection and Estimation of Polyphosphate in Vesicular-Arbuscular Mycorrhizas. New Phytol 80:125–134.

73. Ezawa T, Smith SE, Smith FA. 2002. P metabolism and transport in AM fungi, p. 221–230. In Plant and Soil. Springer.

74. Kanehisa M, Goto S. 2000. KEGG: kyoto encyclopedia of genes and genomes. Nucleic Acids Res 28:27–30.

75. Hastings J, Owen G, Dekker A, Ennis M, Kale N, Muthukrishnan V, Turner S, Swainston N, Mendes P, Steinbeck C. 2016. ChEBI in 2016: Improved services and an expanding collection of metabolites. Nucleic Acids Res 44:D1214–D1219.

76. Lieven C, Beber ME, Olivier BG, Bergmann FT, Ataman M, Babaei P, Bartell JA, Blank LM, Chauhan S, Correia K, Diener C, Dramp A, Ebert BE, Edirisinghe JN, Faria P, Feist AM, Fengos G, T Fleming RM, Garcamp B, Hatzimanikatis V, Helvoirt W, Henry CS, Hermjakob H, Herrgamp MJ, Kaafarani A, Uk Kim H, King Z, Klamt S, Klipp E, Koehorst JJ, Kamp M, Lakshmanan M, Lee D-Y, Yup Lee S, Lee S, Lewis NE, Liu F, Ma H, Machado D, Mahadevan R, Maia P, Mardinoglu A, Medlock GL, Monk JM, Nielsen J, Keld Nielsen L, Nogales J, Nookaew I, Palsson BO, Papin JA, Patil KR, Poolman M, Price ND, Resendis-Antonio O, Richelle A, Rocha I, Samp J, Schaap PJ, Malik Sheriff RS, Shoaie S, Sonnenschein N, Teusink B, Vilaamp P, Olav Vik J, H Wodke JA, Xavier JC, Yuan Q, Zakhartsev M, Zhang C. 2020. MEMOTE for standardized genome-scale metabolic model testing. Nat Biotechnol 38:272–276.

77. Lu H, Li F, Sánchez BJ, Zhu Z, Li G, Domenzain I, Marcišauskas S, Anton PM, Lappa D, Lieven C, Beber ME, Sonnenschein N, Kerkhoven EJ, Nielsen J. 2019. A consensus *S. cerevisiae* metabolic model Yeast8 and its ecosystem for comprehensively probing cellular metabolism. Nat Commun 10:3586.

78. Orth JD, Thiele I, Palsson BO. 2010. What is flux balance analysis? Nat Biotechnol 28:245–248.

79. Savinell JM, Palsson BO. 1992. Optimal selection of metabolic fluxes for in vivo measurement. I. Development of mathematical methods. J Theor Biol 155:201–214.

80. Mahadevan R, Schilling CH. 2003. The effects of alternate optimal solutions in constraint-based genome-scale metabolic models. Metab Eng 5:264–276.

81. Bécard G, Fortin JA. 1988. Early events of vesicular–arbuscular mycorrhiza formation on Ri T-DNA transformed roots. New Phytol 108:211–218.

82. Kameoka H, Tsutsui I, Saito K, Kikuchi Y, Handa Y, Ezawa T, Hayashi H, Kawaguchi M, Akiyama K. 2019. Stimulation of asymbiotic sporulation in arbuscular mycorrhizal fungi by fatty acids. Nat Microbiol. Nature Publishing Group.

83. Pfeffer PE, Douds DD, Bécard G, Shachar-Hill Y. 1999. Carbon Uptake and the Metabolism and Transport of Lipids in an Arbuscular Mycorrhiza. Plant Physiol 120:587–598.

84. Adadi R, Volkmer B, Milo R, Heinemann M, Shlomi T. 2012. Prediction of microbial growth rate versus biomass yield by a metabolic network with kinetic parameters. PLoS Comput Biol 8:e1002575.

85. Bekiaris PS, Klamt S. 2020. Automatic construction of metabolic models with enzyme constraints. BMC Bioinformatics 21:19.

86. Nilsson A, Nielsen J, Palsson BO. 2017. Metabolic Models of Protein Allocation Call for the Kinetome. Cell Syst 5:538–541.

87. Beg QK, Vazquez A, Ernst J, De Menezes MA, Bar-Joseph Z, Barabási AL, Oltvai ZN. 2007. Intracellular crowding defines the mode and sequence of substrate uptake by *Escherichia coli* and constrains its metabolic activity. PNAS 104:12663–12668.

88. Goelzer A, Fromion V, Scorletti G. 2011. Cell design in bacteria as a convex optimization problem. Automatica 47:1210–1218.

89. Li JCH. 2016. Effect size measures in a two-independent-samples case with nonnormal and nonhomogeneous data. Behav Res Methods 48:1560–1574.

90. Perez-Garcia O, Lear G, Singhal N. 2016. Metabolic Network Modeling of Microbial Interactions in Natural and Engineered Environmental Systems. Front Microbiol 7:673.

91. MATLAB. 2017. version 9.3.0 (R2017b). The MathWorks Inc., Natick, Massachusetts.

92. Heirendt L, Arreckx S, Pfau T, Mendoza SN, Richelle A, Heinken A, Haraldsdóttir HS, Wachowiak J, Keating SM, Vlasov V, Magnusdóttir S, Ng CY, Preciat G, Žagare A, Chan SHJ, Aurich MK, Clancy CM, Modamio J, Sauls JT, Noronha A, Bordbar A, Cousins B, El Assal DC, Valcarcel L V., Apaolaza I, Ghaderi S, Ahookhosh M, Ben Guebila M, Kostromins A, Sompairac N, Le HM, Ma D, Sun Y, Wang L, Yurkovich JT, Oliveira MAP, Vuong PT, El Assal LP, Kuperstein I, Zinovyev A, Hinton HS, Bryant WA, Aragón Artacho FJ, Planes FJ, Stalidzans E, Maass A, Vempala S, Hucka M, Saunders MA, Maranas CD, Lewis NE, Sauter T, Palsson BØ, Thiele I, Fleming RMT. 2019. Creation and analysis of biochemical constraint-based models using the COBRA Toolbox v.3.0. Nat Protoc 14:639–702.

93. King ZA, Lu J, Dräger A, Miller P, Federowicz S, Lerman JA, Ebrahim A, Palsson BO, Lewis NE. 2016. BiGG Models: A platform for integrating, standardizing and sharing genome-scale models. Nucleic Acids Res 44:D515–D522.

94. Caspi R, Billington R, Keseler IM, Kothari A, Krummenacker M, Midford PE, Ong WK, Paley S, Subhraveti P, Karp PD. 2020. The MetaCyc database of metabolic pathways and enzymes-a 2019 update. Nucleic Acids Res 48:D455–D453.

95. Moretti S, Martin O, Van Du Tran T, Bridge A, Morgat A, Pagni M. 2016. MetaNetX/MNXref - Reconciliation of metabolites and biochemical reactions to bring together genome-scale metabolic networks. Nucleic Acids Res 44:D523–D526.

96. Kim S, Chen J, Cheng T, Gindulyte A, He J, He S, Li Q, Shoemaker BA, Thiessen PA, Yu B, Zaslavsky L, Zhang J, Bolton EE. 2019. PubChem 2019 update: Improved access to chemical data. Nucleic Acids Res 47:D1102–D1109.

97. Jolicoeur M, Germette S, Gaudette M, Perrier M, Bécard G. 1998. Intracellular pH in Arbuscular Mycorrhizal Fungi: A Symbiotic Physiological Marker. Plant Physiol 116:1279–1288.

98. Belmondo S, Fiorilli V, Pérez-Tienda J, Ferrol N, Marmeisse R, Lanfranco L. 2014. A dipeptide transporter from the arbuscular mycorrhizal fungus *Rhizophagus irregularis* is upregulated in the intraradical phase. Front Plant Sci 5:436.

99. Chan SHJ, Cai J, Wang L, Simons-Senftle MN, Maranas CD. 2017. Standardizing biomass reactions and ensuring complete mass balance in genome-scale metabolic models. Bioinformatics 33:3603–3609.

100. Sánchez BJ, Li F, Kerkhoven EJ, Nielsen J. 2019. SLIMEr: Probing flexibility of lipid metabolism in yeast with an improved constraint-based modeling framework. BMC Syst Biol 13:1–9.

101. Maranas CD, Zomorrodi AR. 2016. Thermodynamic Analysis of Metabolic Networks, p. 107–117. In Optimization Methods in Metabolic Networks. John Wiley & Sons, Ltd.

102. Bolger AM, Lohse M, Usadel B. 2014. Trimmomatic: a flexible trimmer for Illumina sequence data. Bioinformatics 30:2114–2120.

103. Dobin A, Davis CA, Schlesinger F, Drenkow J, Zaleski C, Jha S, Batut P, Chaisson M, Gingeras TR. 2013. STAR: ultrafast universal RNA-seq aligner. Bioinformatics 29:15–21.

104. Anders S, Pyl PT, Huber W. 2015. HTSeq--a Python framework to work with high-throughput sequencing data. Bioinformatics 31:166–169.

105. Altschul SF, Gish W, Miller W, Myers EW, Lipman DJ. 1990. Basic local alignment search tool. J Mol Biol 215:403–410.

106. Camacho C, Coulouris G, Avagyan V, Ma N, Papadopoulos J, Bealer K, Madden TL. 2009. BLAST+: architecture and applications. BMC Bioinformatics 10:421.

107. Placzek S, Schomburg I, Chang A, Jeske L, Ulbrich M, Tillack J, Schomburg D. 2017. BRENDA in 2017: new perspectives and new tools in BRENDA. Nucleic Acids Res 45:D380–D388.

108. Wittig U, Rey M, Weidemann A, Kania R, Müller W. 2018. SABIO-RK: An updated resource for manually curated biochemical reaction kinetics. Nucleic Acids Res 46:D656–D660.

109. The UniProt Consortium. 2017. UniProt: the universal protein knowledgebase. Nucleic Acids Res 45:D158–D169.

110. Meijer MMC, Boonstra J, Verkleij AJ, Theo Verrips C. 1996. Kinetic analysis of hexose uptake in *Saccharomyces cerevisiae* cultivated in continuous culture. Biochim Biophys Acta - Bioenerg 1277:209–216.

111. Reifenberger E, Boles E, Ciriacy M. 1997. Kinetic Characterization of Individual Hexose Transporters of *Saccharomyces cerevisiae* and their Relation to the Triggering Mechanisms of Glucose Repression. Eur J Biochem 245:324–333.

